## Supplementary figures and images for "DNA variants affecting the expression of numerous genes in *trans* have diverse mechanisms of action and evolutionary histories"

### Supplemental Figure 1

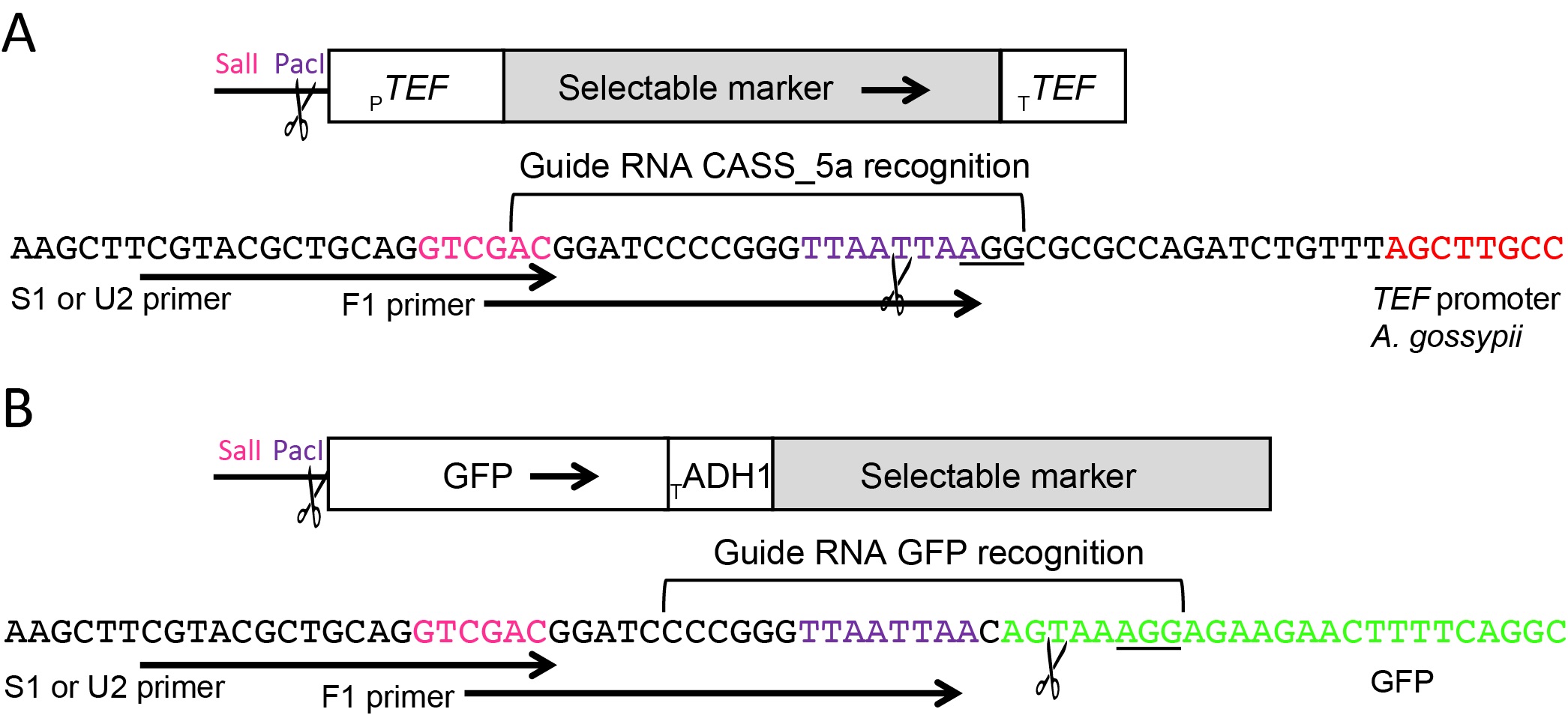

### Supplemental Figure 2

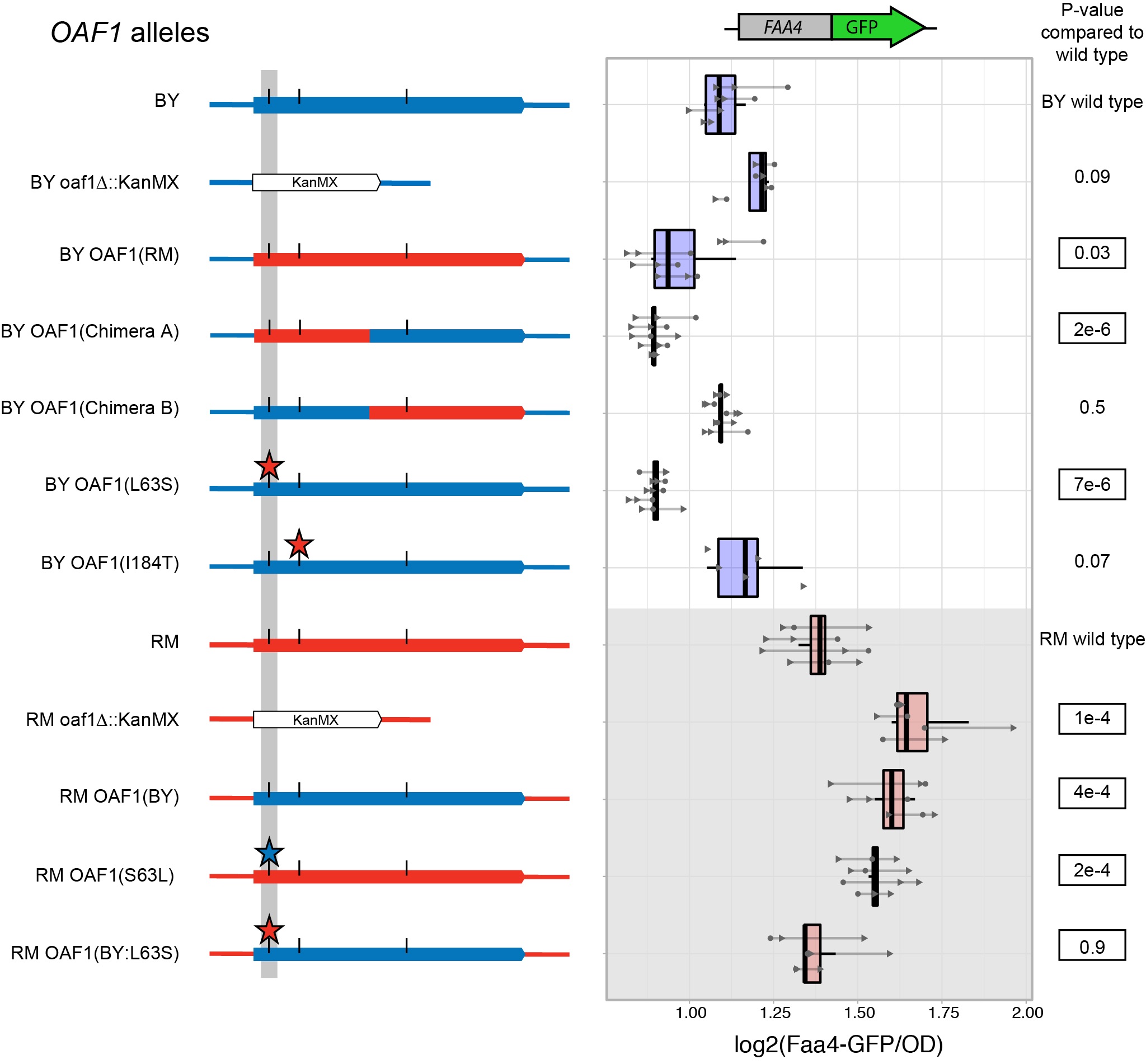

### Supplemental Figure 3

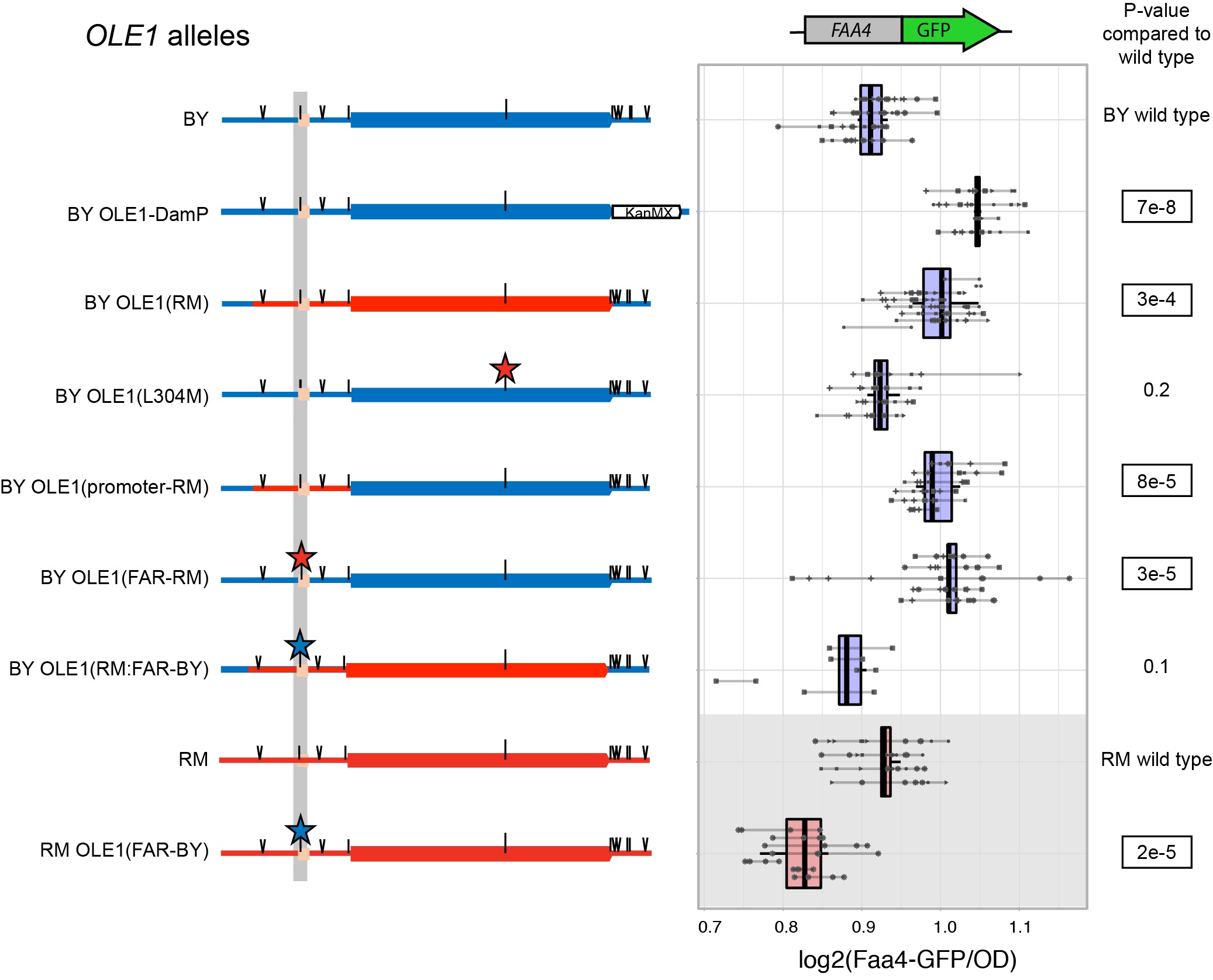

### Supplemental Figure 4

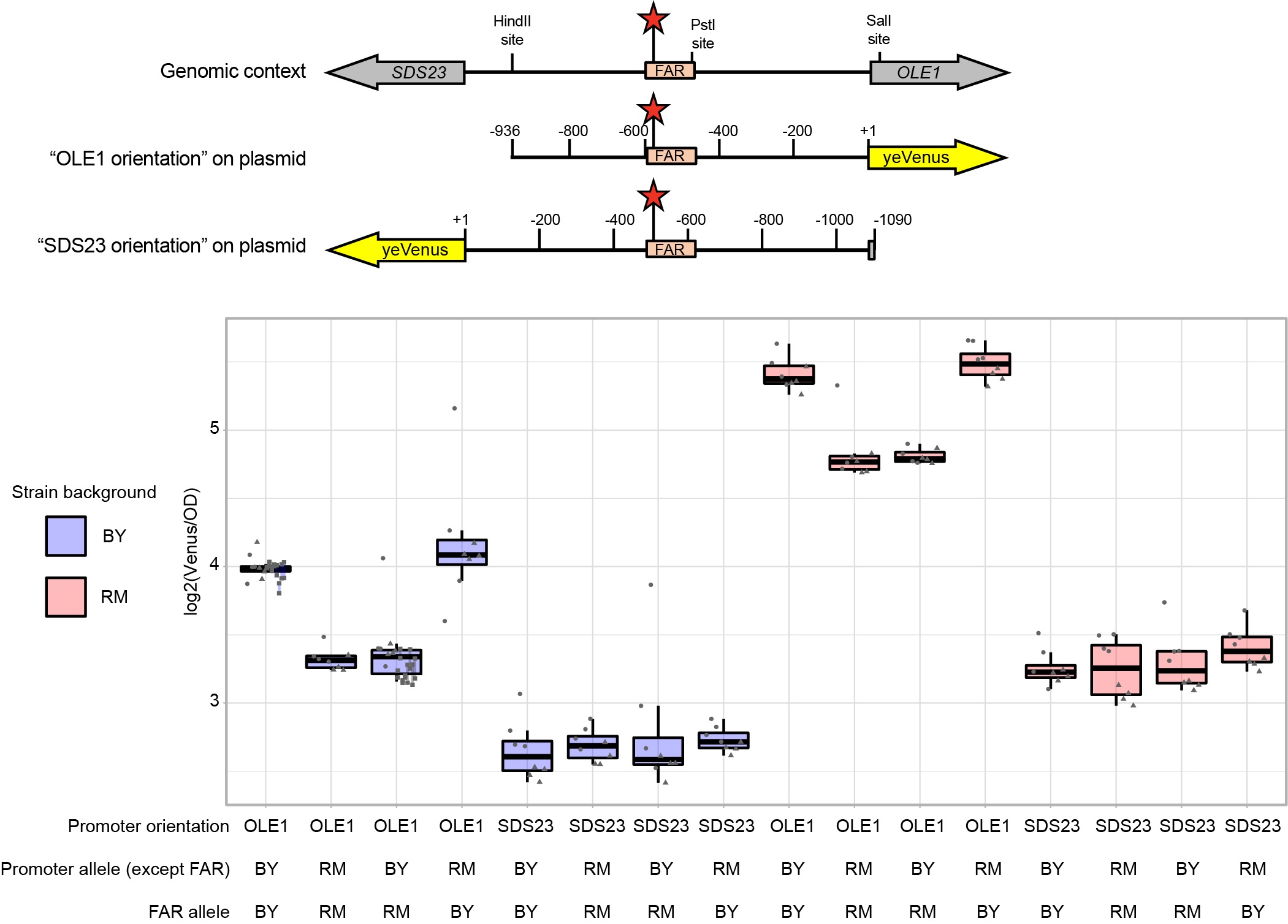

### Supplemental Figure 5

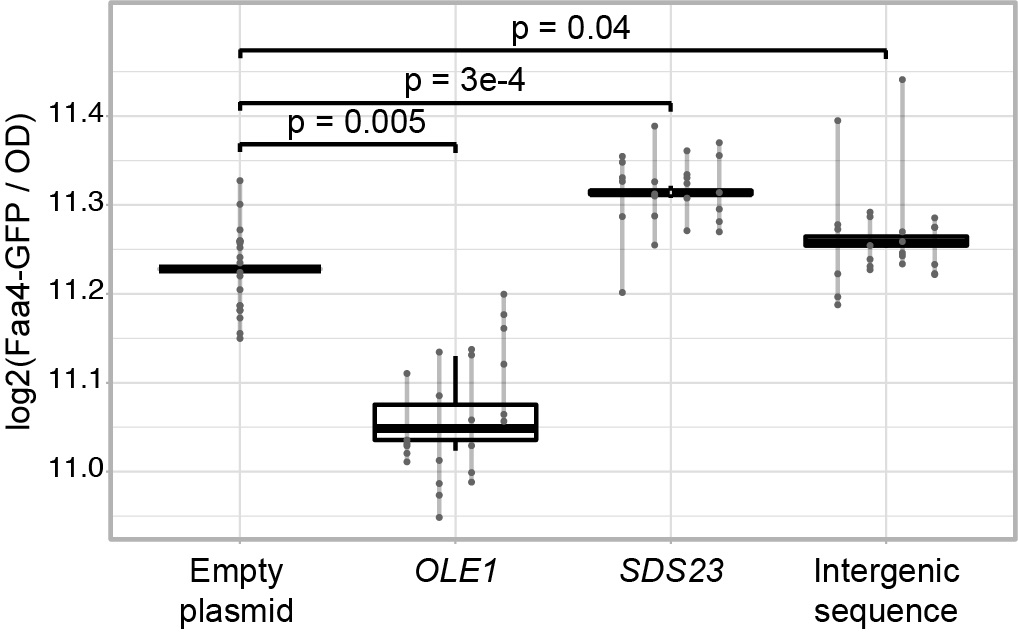

### Supplemental Figure 6

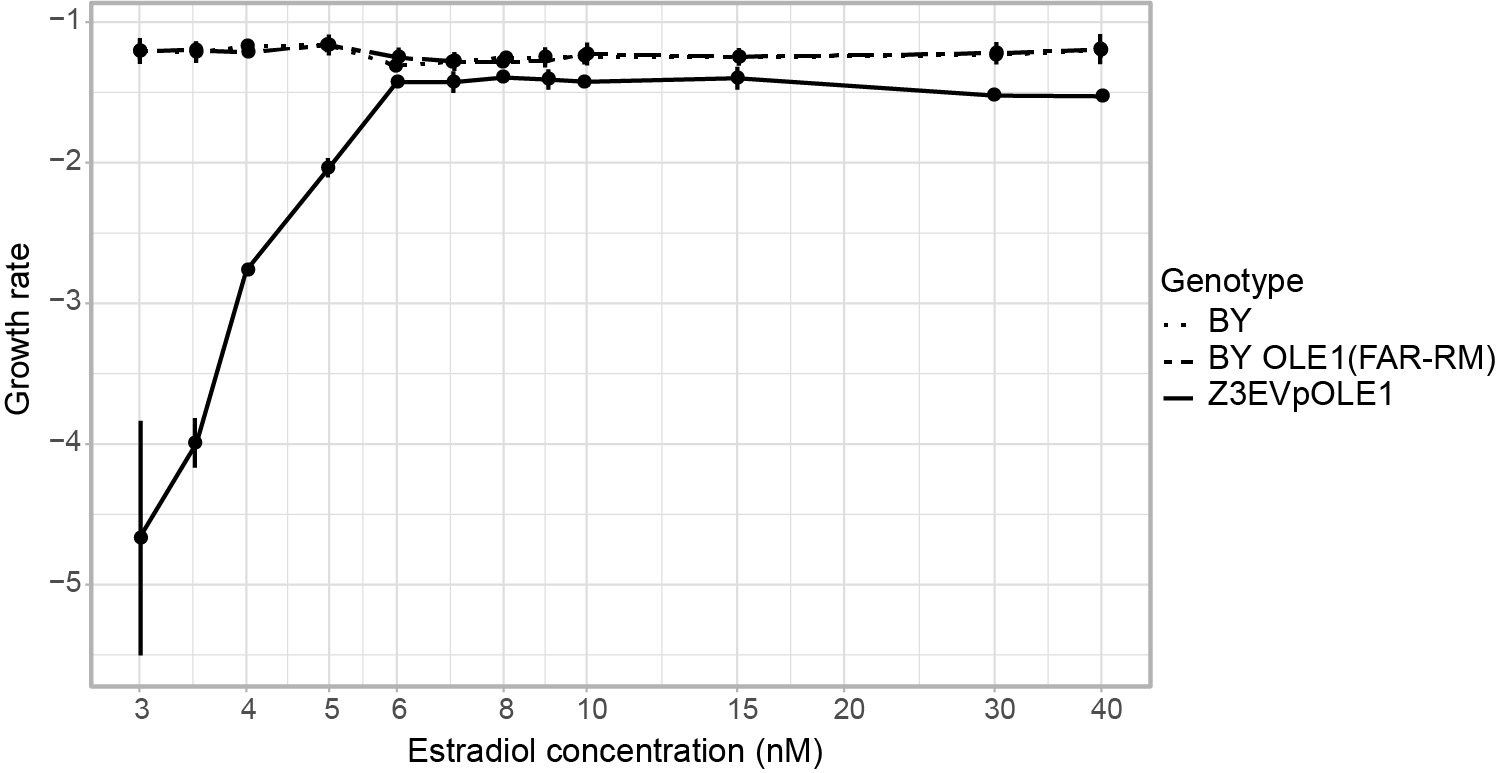

### Supplemental Figure 7

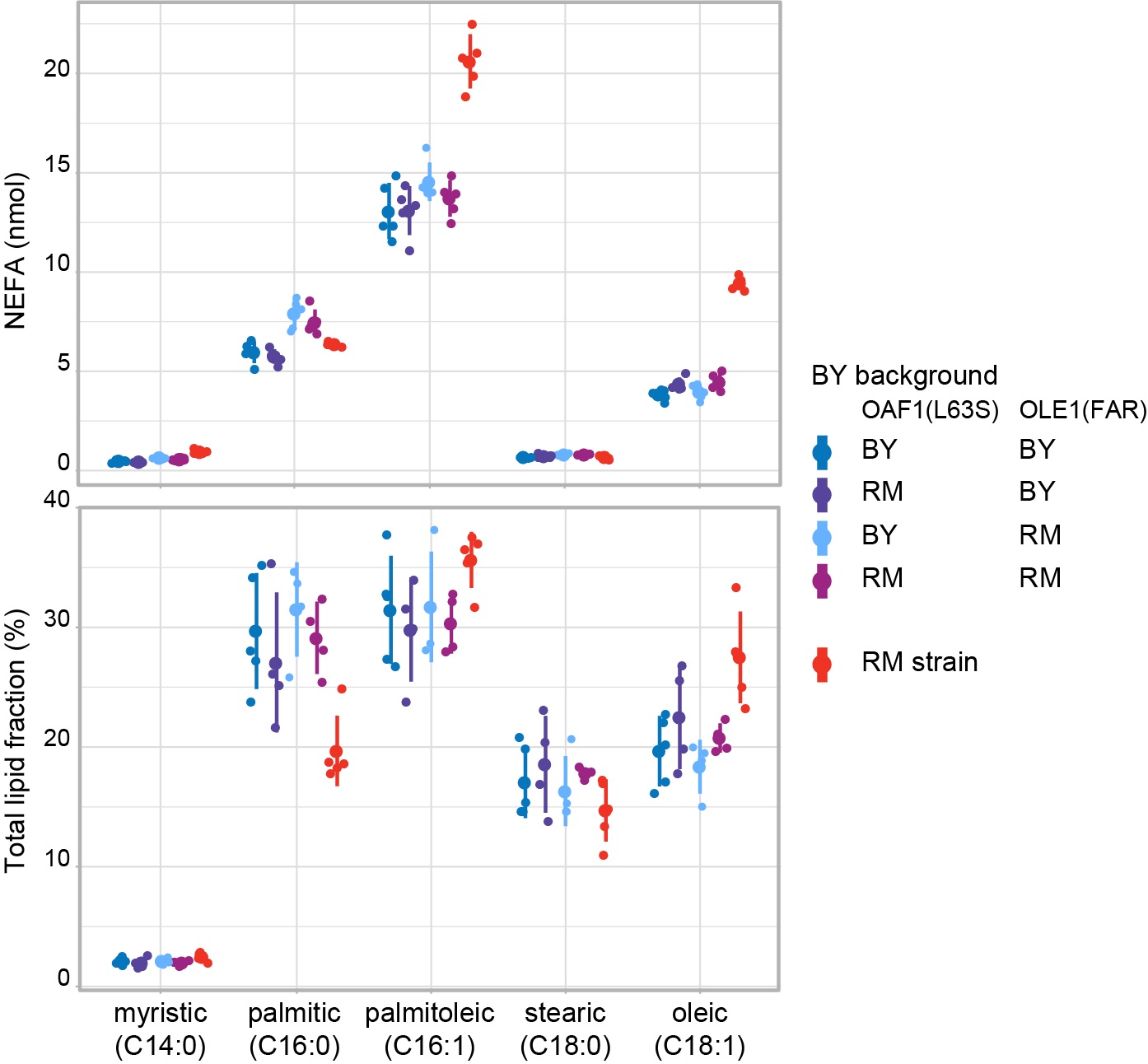
